# The immediate cellular response to whole-genome doubling is conserved across polyploid contexts

**DOI:** 10.64898/2026.07.07.736946

**Authors:** Cindy C. Geerlings, Gabriella S. Darmasaputra, Christa Jordan Ortiz, Susana M. Chuva de Sousa Lopes, Hans Clevers, Matilde Galli

## Abstract

Polyploid cells, which contain more than two copies of the genome, are widely present across plants and animals, where they are often found in tissues with high biosynthetic and metabolic demands, such as the mammalian liver and placenta. While somatic polyploidy is frequently associated with increased cell growth and biosynthetic capacity, unscheduled polyploidization in cell types that are not normally programmed to become polyploid is often linked to reduced cellular fitness and genome instability. To understand whether these divergent outcomes stem from distinct immediate cellular responses to increased ploidy, we systematically compared the early consequences of polyploidization across naturally occurring and experimentally induced systems. Specifically, we examined physiological polyploid cells in the *Caenorhabditis elegans* intestine and human hepatocyte organoids, alongside unscheduled polyploid human retinal pigment epithelial (RPE1) cells generated through cytokinesis failure. Using quantitative imaging, flow cytometry, and FUCCI-based cell-cycle reporters we measured cell size and protein translation dynamics during G1 in diploid and polyploid cells. Across all systems, we observed a strikingly conserved relationship between ploidy, cell size, and biosynthetic capacity: both cell size and protein translation showed similar scaling patterns after polyploidization, regardless of whether polyploidization occurred as part of normal development or by inducing cytokinesis failure. These findings indicate that the immediate cellular response to increased ploidy is broadly similar across contexts. However, in contrast to unscheduled polyploid RPE1 cells, polyploid human hepatocytes extend their G1 phase, leading to a higher accumulation of proteins before cell-cycle progression. Together, our findings suggest that polyploidization elicits similar growth responses across contexts, and that cell-type specific cell-cycle adaptations may determine whether polyploidy becomes advantageous or deleterious.

## Introduction

Polyploid cells, which contain more than two copies of the genome, are frequently observed in many cell types across multicellular organisms. In mammals, some examples of polyploid cell types include megakaryocytes, hepatocytes and cardiomyocytes, but polyploid cells can be found in many other animal and plant tissues^1,2^. Somatic polyploidy can arise either by cell fusion or by non-canonical cell cycles in which cells replicate their DNA but do not divide into two daughter cells. Two types of non-canonical cell cycles exist: endomitosis and endoreplication. During endomitosis, cells enter M phase but exit prematurely, resulting in either mononucleated or binucleated polyploid cells, whereas during endoreplication cells skip M phase entirely and repeatedly alternate between gap and S phases to increase their DNA content^3^. Importantly, these non-canonical cell cycles occur as part of healthy development and are tightly regulated^3^.

Many polyploid cell types perform specialized functions during development^1^. For example, polyploid mammary epithelial cells support efficient milk production, polyploid cells lining the urothelium have a barrier function, and polyploid intestinal cells in *C. elegans* have important functions to fine-tune gene expression^4–6^. Despite the widespread occurrence of polyploidy and its association with diverse cellular functions, it is not fully understood why cells adopt a polyploid state rather than simply increasing in cell numbers. Furthermore, little is known about the direct molecular and cellular consequences of whole-genome doubling (WGD), for example how cells adjust their gene expression, protein production and cellular growth after becoming polyploid. It is possible that polyploidization confers specific advantages to certain molecular processes such as protein translation, however, this has not been systematically addressed.

One common feature of polyploid cells is that they are bigger than diploid cells, a phenomenon that has been observed across multiple organisms and cell types^1^. For example, cell size scales with DNA content in several mammalian cell types, including hepatocytes, cardiomyocytes, megakaryocytes, and neurons^7–11^. However, most studies supporting this association are correlative, and direct evidence that polyploidy drives an increase in cell and tissue size has been demonstrated only in a few systems. In *C. elegans*, experimentally increasing or decreasing polyploidization in the hypodermis leads to corresponding increases or decreases in body size, respectively^12^. Similarly, in *Drosophila melanogaster* polyploidization has an important function in the regulation of cell and tissue size, as blocking DNA replication reduces cell and organ size in subperineurial glial cells (SPGs), salivary glands, and hair bristles; conversely, increasing ploidy levels results in larger SPGs and brain lobes^13–15^. Together, these findings demonstrate that polyploidy can directly drive increases in cell and tissue size across diverse organisms.

Although somatic polyploidy appears to be beneficial in many cell types, experimentally inducing polyploidization often leads to detrimental cellular consequences. Studies in yeast have shown that increased ploidy can result in genetic instability and slower growth^16–20^. Similar observations have been made in animal cells, where induction of tetraploidy through cytokinesis failure frequently triggers cell-cycle arrest or apoptosis^21,22^. One proposed explanation for the reduced fitness of induced polyploid cells is an imbalance in protein synthesis, which insufficiently scales with the increased ploidy^23,24^. Supporting this idea, RNA and protein content have been shown to increase sublinearly with ploidy in budding yeast^23^. Furthermore, *in vitro* induction and selection of tetraploidy in human colorectal cancer cells results in a downregulation of ribosomal proteins and reduced translation^23^. Also in non-cancerous human cells, induction of polyploidy results in an insufficient accumulation of replication factors during G1, causing DNA damage in the subsequent S phase^24^. Together, these findings suggest that cells that become polyploid following cytokinesis failure may experience reduced fitness due to imbalances between their genomic content, cell growth, and protein synthesis. In contrast, naturally occurring polyploid cells are typically associated with enhanced growth and increased protein production. This raises the possibility that the consequences of polyploidization are context-dependent and could vary depending on the cell type or the mechanism by which polyploidy arises.

To understand why polyploidization leads to different outcomes in naturally occurring versus induced polyploid cells, we systematically compared cell size and protein synthesis across diploid and polyploid cells in three different contexts: two naturally occurring polyploid cell types, *C. elegans* intestinal cells and human hepatocytes, and in human RPE1 cells in which polyploidy was induced through cytokinesis failure. Strikingly, we find that both cell size and nascent protein synthesis scale similarly with ploidy in both naturally occurring and induced polyploid cells. However, we find that human hepatocytes lengthen their G1 phase after polyploidization, allowing sufficient biosynthetic accumulation before DNA replication, whereas experimentally induced polyploid cells fail to do this. Together, our results show that the immediate effects of polyploidization on growth and protein production are largely similar across cell types, and that the reduced fitness observed in induced polyploid cells likely arises from a failure to extend the cell cycle to accommodate the increased biosynthetic demands of polyploid growth.

## Results

### Cell size scales linearly with ploidy in both naturally occurring and induced polyploid cells

To understand how polyploidization affects cellular functions and whether there are different responses in naturally occurring polyploid cells versus cells that undergo an unscheduled polyploidization event, we first investigated how cell size scales with ploidy in these different contexts. As model systems for naturally occurring polyploidy, we used the *C. elegans* intestine and fetal-derived human hepatocyte organoids, which both have been previously shown to undergo programmed polyploidization^25–28^. The *C. elegans* intestine is composed of 20 cells that become polyploid during larval development by performing two types of non-canonical cycles: an endomitosis cell cycle at the end of the first larval stage (L1) which results in binucleated cells, followed by endoreplication cycles at the end of each larval stage (L1 to L4) (**Figure 1A**). To investigate how polyploidy influences cell growth, we quantified changes in intestinal cell size over developmental time and compared them to body-wall muscle cells, which are postmitotic and remain diploid throughout larval development. We used a *C. elegans* strain expressing GFP::PH and mCherry::PH to label the membranes of intestinal and body-wall muscle cells, respectively. DNA content was verified by propidium iodide (PI) staining at the beginning of each larval stage (**Figure S1A**), confirming the expected ploidy levels. We then performed live imaging at matched timepoints across larval stages and quantified cell volumes. Intestinal cell volume increases during larval development in parallel with rising ploidy levels (**Figure 1B**). Notably, body-wall muscle cells, which remain diploid, also exhibit similar increases in cell size over the same developmental period (**Figure 1B, C**), suggesting that growth is not solely driven by changes in ploidy.

**Figure 1.**
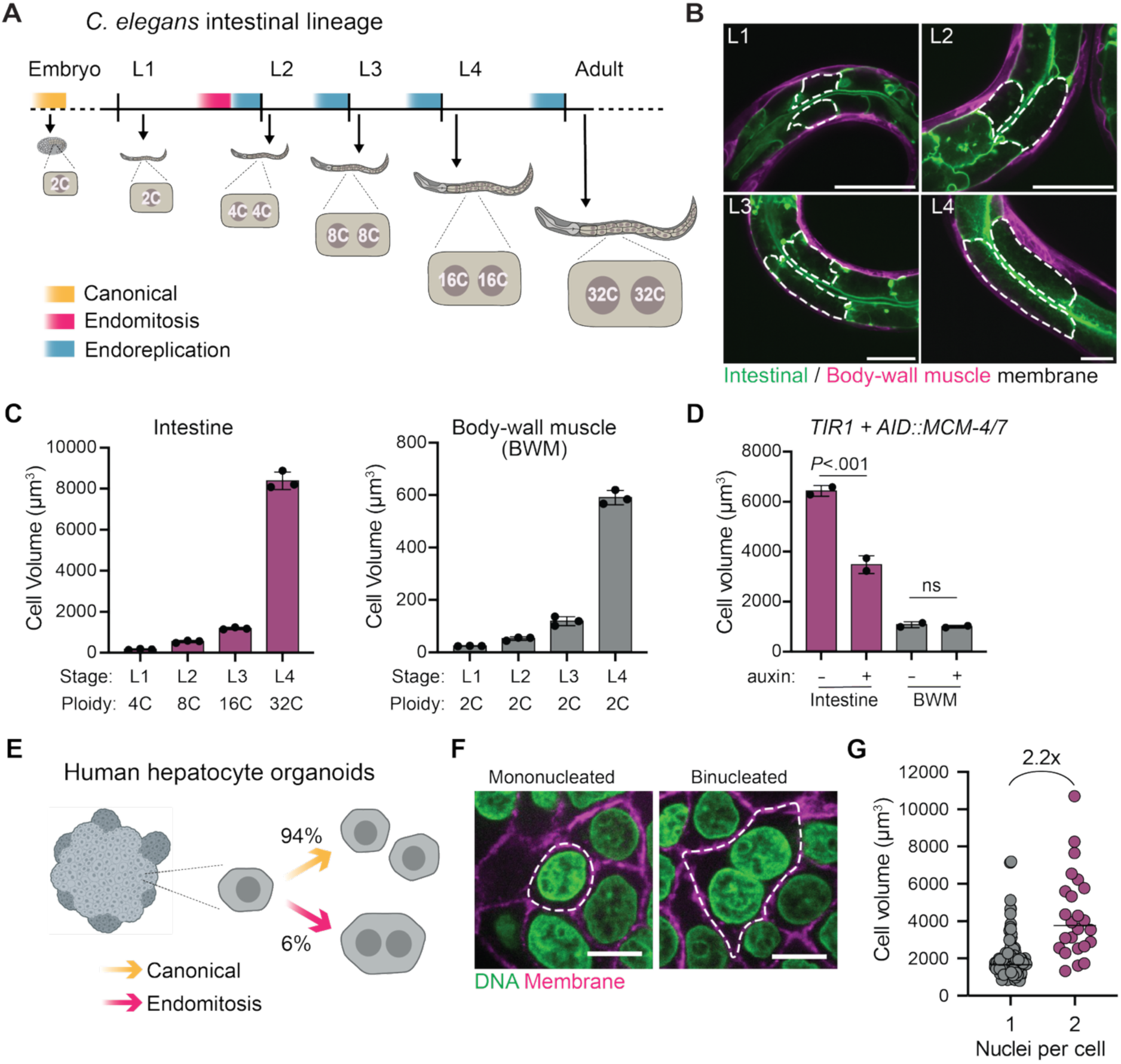
Cell size scales linearly with ploidy in naturally occurring polyploid cells. **(A)** Schematic of cell-cycle transitions and ploidy increases in Int3V/D intestinal cells during *C. elegans* larval development. During larval stages L1 to L4, the intestinal lineage undergoes non-canonical cell cycles (endomitosis and endoreplication), resulting in the duplication of the genome at the end of each larval stage, giving rise to adult intestinal cells with a cellular ploidy of 64C DNA content, distributed over two nuclei. **(B)** Representative images of *C. elegans* strain expressing GFP-PH (intestinal membrane, green) and mCherry-PH (body-wall muscle membrane, magenta) during larval stages. Dashed lines indicate cell outlines of intestinal Int3V/D cells. Scale bar is 20 μm. **(C)** Quantification of Int3V/C cell volumes (left; purple) and body-wall muscle cell volumes (right; gray) in all four larval stages. Columns indicate mean ± SD (N = 3). **(D)** Quantification of intestinal cell volume (purple bars, left) of intestinal ring 3 and body-wall muscle cell volume (gray bars, right) in L4 stage *Pges-1::TIR1; AID::mcm-4; AID::mcm-7* animals grown with auxin (+) or without auxin (-). Dots show average cell volumes in individual replicates and columns depict mean ± SD (N = 2, ns = not significant, two-tailed Student’s *t* test). **(E)** Schematic of hepatocyte cell cycles in Hep-Orgs. **(F)** Representative images of mononucleated (left) and binucleated (right) cells in Hep-Orgs expressing E-cadherin-tdTomato (magenta) and stained with DAPI (green). Images show a single plane from a 3D organoid grown in BME gel matrix. Dashed white lines indicate cell outline. Scale bars represent 10 µm. **(G)** Cell volume measurements of mononucleated (gray) and binucleated (purple) hepatocytes. Each point represents one cell (n = 151 mononucleated cells and n = 24 binucleated cells from N = 3 independent experiments). Black lines indicate median values, fold change between medians is indicated above.

To disentangle the contributions of polyploidization and developmental progression to intestinal cell growth, we next examined how and to what extent polyploidy directly influences cell size. To manipulate intestinal ploidy in a tissue-specific and inducible manner, we made use of the auxin-inducible degron (AID) system, in which a target protein is tagged with a degron that promotes its degradation upon expression of the auxin receptor TIR1 and exposure to auxin^29–31^. We generated a *C. elegans* strain in which two subunits of the minichromosome maintenance (MCM) complex, MCM-4 and MCM-7, which are essential for DNA replication^32^, were endogenously tagged with AID degrons. Crossing this strain to animals expressing intestinal driven Tir1 and membrane markers for intestinal and body-wall muscle cells, allowed us to determine how intestinal polyploidization affects cell growth.

We first verified that the degradation of MCM-4 and MCM-7 was sufficient to inhibit replication and thereby reduce intestinal ploidy. Synchronized L1 larvae were grown on either control or auxin plates until the L4 stage, after which DNA content was measured by propidium iodide (PI). Intestinal ploidy in auxin-treated animals was reduced to 16C ± 1.18 compared to 28C ± 0.89 in control animals (mean ± SEM, N = 2, **Figure S1B**). We next performed live imaging of animals with and without reduced intestinal ploidy and quantified cell volumes of intestinal and body-wall muscle cells. Animals grown on auxin showed reduced intestinal cell volume compared to controls (**Figure 1D, S1C**). Notably, the reduction in intestinal cell volume was proportional to the decrease in ploidy, which were both halved in auxin-treated animals. In contrast, body-wall muscle cell volumes showed no statistically significant reduction in the auxin-treated condition. These results indicate that polyploidization plays an important role in determining intestinal cell size, and that increased ploidy levels are important to sustain an increased intestinal cell size in *C. elegans*.

To understand if cell size scales similarly in polyploid mammalian cells, we made use of a fetal-derived human hepatocyte organoid (Hep-Org) system, in which we previously showed that binucleated cells arise through endomitosis and make up around six percent of the cell population^28^ (**Figure 1E**). When Hep-Orgs are grown three-dimensionally in a basement membrane extract (BME) gel matrix, binucleated cells can be found throughout the organoid (**Figure 1F**). Since it is challenging to quantify DNA content reliably in 3D organoids, as fluorescence intensity depends on the focal plane and nuclei at different depths cannot be directly compared, we first compared cell volumes of mono- and binucleated hepatocytes. We used a Hep-Org line expressing GFP-NLS and endogenously-tagged E-cadherin-tdTomato to label nuclei and cell membranes, respectively, and quantified cell volumes using a prismatoid model based on the areas of the top, middle, and bottom planes (**Figure S1D**). We found that the median cell volume of binucleated cells is 2.2-fold larger than the volume of mononucleated cells (**Figure 1G**). These results suggest that cell size also scales linearly with ploidy in naturally occurring mammalian polyploid hepatocytes.

Because cells grow throughout the cell cycle and polyploid hepatocytes have been shown to cycle slower^33^, the observed differences in cell volume between mononucleated and binucleated cells could be explained by differences in cell age rather than differences in ploidy. To enable direct comparison of diploid and polyploid hepatocyte volumes from their birth throughout G1 phase, we integrated a fluorescent ubiquitination-based cell-cycle indicator (FUCCI) in Hep-Orgs. The FUCCI system labels cells with an orange fluorescent protein in G1, early S phase and G0, and a green fluorescent protein during S, G2 and early M phase, allowing determination of the cell-cycle phase of each cell^34^ (**Figure 2A, B**). Importantly, the G1 marker (Cdt1) accumulates progressively during G1, allowing its fluorescence to be used as a proxy for progression through G1 as was shown previously^35^. To directly quantify DNA content in single cells while controlling for the cell-cycle stage, we performed calibrated flow cytometry (see **Methods** and **Figure S2, S3A**). This approach enabled simultaneous measurement of cell size, DNA content, and cell-cycle phase using Hep-Orgs expressing the FUCCI reporter. The FUCCI reporter was used to gate cells in G1 phase, which were then gated for 2N and 4N ploidy based on DNA staining (**Figure S2, S3B**). The dynamic range of the Cdt1 signal was broadly similar between 2N and 4N cells (**Figure S3C**). Consistent with our previous work, approximately 6% of the dissociated organoid cells displayed a 4N ploidy^28^ (**Figure S3B**). Analysis of hepatocyte size distributions across increasing Cdt1 signals revealed that 4N hepatocytes are on average 2.00 ± 0.08-fold larger than 2N hepatocytes (mean ± SEM, N = 4, **Figure 2C**). Notably however, the extent of cell growth throughout G1 was very similar between 2N and 4N populations (**Figure 2C**). These findings validate our imaging-based measurements and demonstrate that cell size scales proportionally with ploidy in human hepatocytes. Together with our observations in *C. elegans*, these results indicate that proportional scaling of cell size with genome content is a conserved feature of naturally occurring polyploid cell types.

**Figure 2.**
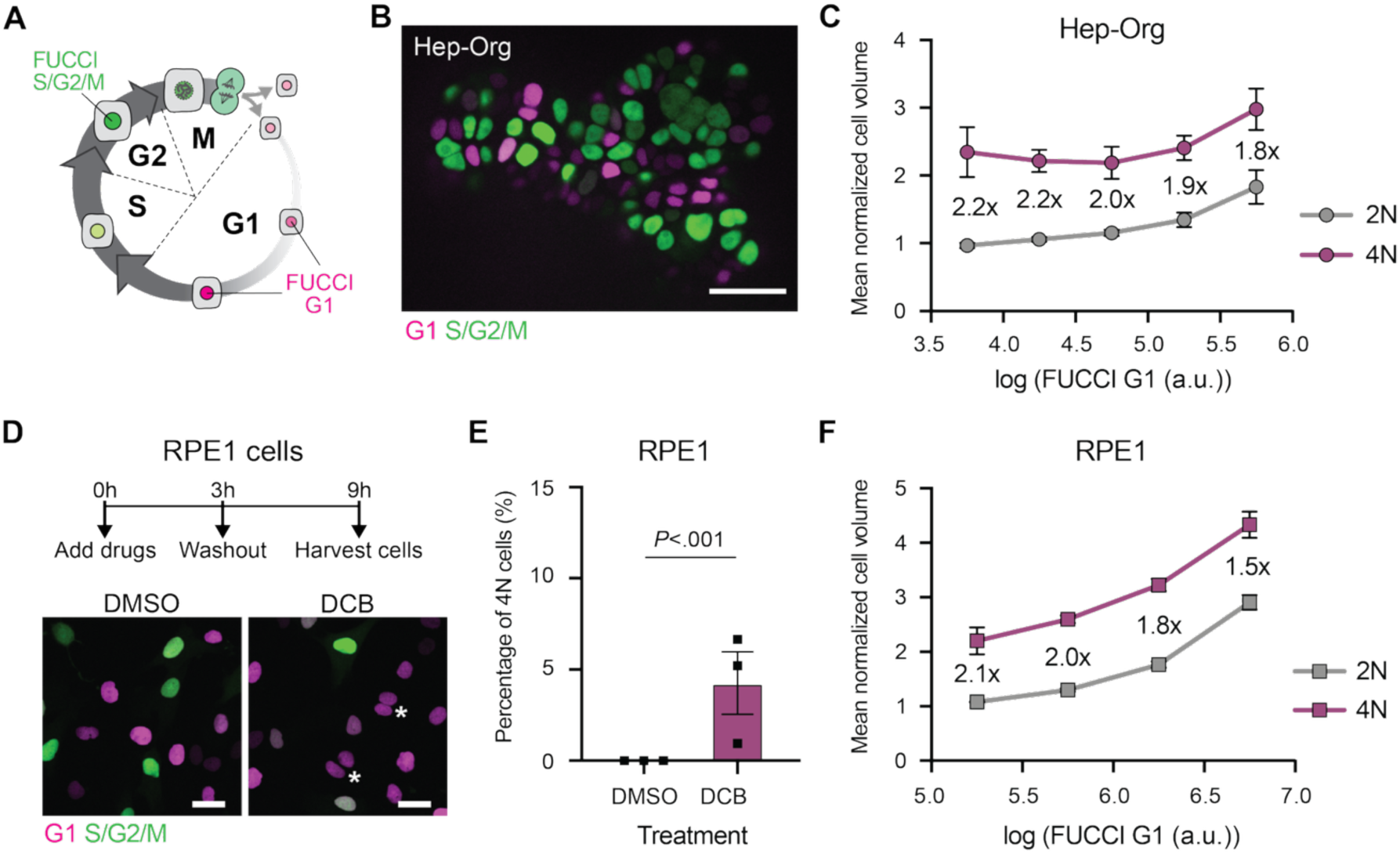
Cell sizes scale similarly with ploidy in both naturally occurring and induced mammalian polyploid cells. **(A**) Schematic of the fluorescent ubiquitination-based cell-cycle indicator (FUCCI) system used to distinguish cell-cycle phases. Cdt1 (magenta) marks cells in G1 phase and Geminin (green) marks cells in S/G2/M phases. **(B)** Representative image of a Hep-Org expressing FUCCI markers (Cdt1, magenta; Geminin, green). Image shows a single z-plane in the middle of a 3D organoid. Scale bar represents 50 µm. **(C)** Normalized volumes of Hep-Org cells, measured by calibrated flow cytometry. G1-phase cells were binned in five equal bins. Mean values from N = 4 independent replicates are shown for 2N (gray, n > 7500 cells per replicate) and 4N (purple, n > 220 cells per replicate) cells. Error bars represent SD. Fold changes are indicated per bin, overall fold change is 2.00 ± 0.08 (mean ± SEM). **(D)** Schematic and representative images of the induction of polyploidy in RPE1 cells. RPE1 cells were treated with DMSO or DCB for 3 hours, after which the drug was washed out, and the newly formed polyploid cells were allowed to progress through G1 phase for 6 hours before harvesting cells for cell volume analysis (h = hours). Images show the nuclear signal of Cdt1 (magenta) and Geminin (green), white asterisks mark induced binucleated polyploid cells and scale bar represents 20 µm. **(E)** Percentage of cells with 4N ploidy upon DMSO or DCB treatment in RPE1 cells. Bar shows the mean and dots show average percentages in individual replicate experiments. Error bars represent SEM (N = 3, n > 10^4^ cells per experiment, \*\*\**P* < 0.001 paired Student’s *t* test). **(F)** Normalized cell volumes of diploid and induced polyploid RPE1 cells. Cell volumes were measured by calibrated flow cytometry and binned in four equal bins based on log (FUCCI G1) signal. Mean normalized cell volume is plotted from N = 3 independent replicate experiments for 2N (gray, n > 95000 cells per replicate) and 4N (purple, n > 1420 cells per replicate) cells. Error bars represent SD. Fold changes are indicated per bin, overall fold change is 1.98 ± 0.02 (mean ± SEM).

Next, we set out to investigate how cell size scales with ploidy in cells that are experimentally induced to become polyploid. Previous studies have reported increased cell size following cytokinesis failure in human cells, consistent with the general association between higher ploidy and larger cell volume^1,23,24,36^. However, these analyses were largely based on population-level averages and did not account for differences in cell-cycle distributions between diploid and polyploid populations^23,24,36^. To determine whether induced polyploidization gives rise to differences in cell growth during G1, we generated polyploid cells in the near-diploid untransformed RPE1 cell line expressing the FUCCI reporter by transient treatment with dihydrocytochalasin B (DCB), which induces cytokinesis failure^37,38^ (**Figure 2D**). Cells were treated with DCB or DMSO for three hours, after which they were allowed to recover for six hours to minimize drug effects and allow investigation of newly formed induced polyploid cells. Using this assay, we observed an average of 4% polyploid cells in G1 after DCB treatment, compared to 0% after DMSO treatment (**Figure 2E**). We then used calibrated flow cytometry to measure cell size in cells with different ploidies. Again, both diploid and induced polyploid cells increased in size during G1 (**Figure 2F**). Surprisingly, induced polyploid cells increased in size to 1.98 ± 0.02 times that of diploid cells (mean ± SEM, N = 3), which was a similar proportional scaling of cell size with ploidy as observed in hepatocytes (**Figure 2C, F**). Together, these results suggest that the immediate consequences of WGD on cell growth are similar between naturally occurring and induced mammalian polyploid cells.

### Nascent protein synthesis increases similarly with ploidy both in naturally occurring and induced polyploid cells

Our analyses of cell volumes in naturally occurring and induced polyploid cells show that cell size scales proportionally with ploidy in both types of polyploid cells, suggesting that cell growth is not immediately affected after a WGD event. However, previous studies have demonstrated that induced polyploidy impairs cellular fitness, including reduced protein synthesis in both yeast and human cells^16,20,23,24^. This raises the possibility that, despite scaling of cell size with ploidy, biosynthetic capacity fails to scale in induced polyploid cells. To compare protein synthesis levels in diploid and induced polyploid RPE1 cells we used O-propargyl-puromycin (OPP), an analog of puromycin, to visualize protein translation^39^. When OPP is added to cells, it is incorporated into newly synthesized proteins, causing premature termination of the polypeptide chain (**Figure 3A**). The resulting OPP-labeled proteins can then be fixed and labeled with a fluorescent dye by click chemistry. To measure the effect of polyploidization on protein synthesis, we performed OPP assays in RPE1 cells treated with DMSO or DCB for two to three hours and allowed to recover for four hours, after which cells were analyzed by flow cytometry to measure Cdt1, Geminin, Hoechst and OPP intensities. Comparison of OPP signals throughout G1 in diploid and polyploid RPE1 cells revealed that induced polyploid cells produce 1.64 ± 0.03 more protein in G1 than their diploid counterparts (mean ± SEM, N = 3, **Figure 3B**). These measurements are in line with the sublinear scaling of protein content with ploidy that has been observed in yeast, which was shown to be due to reduced translation^23^. We thus conclude that despite an increase in protein translation in induced polyploid cells relative to diploid cells, induced polyploidization does not lead to a proportional scaling of protein synthesis with ploidy.

**Figure 3.**
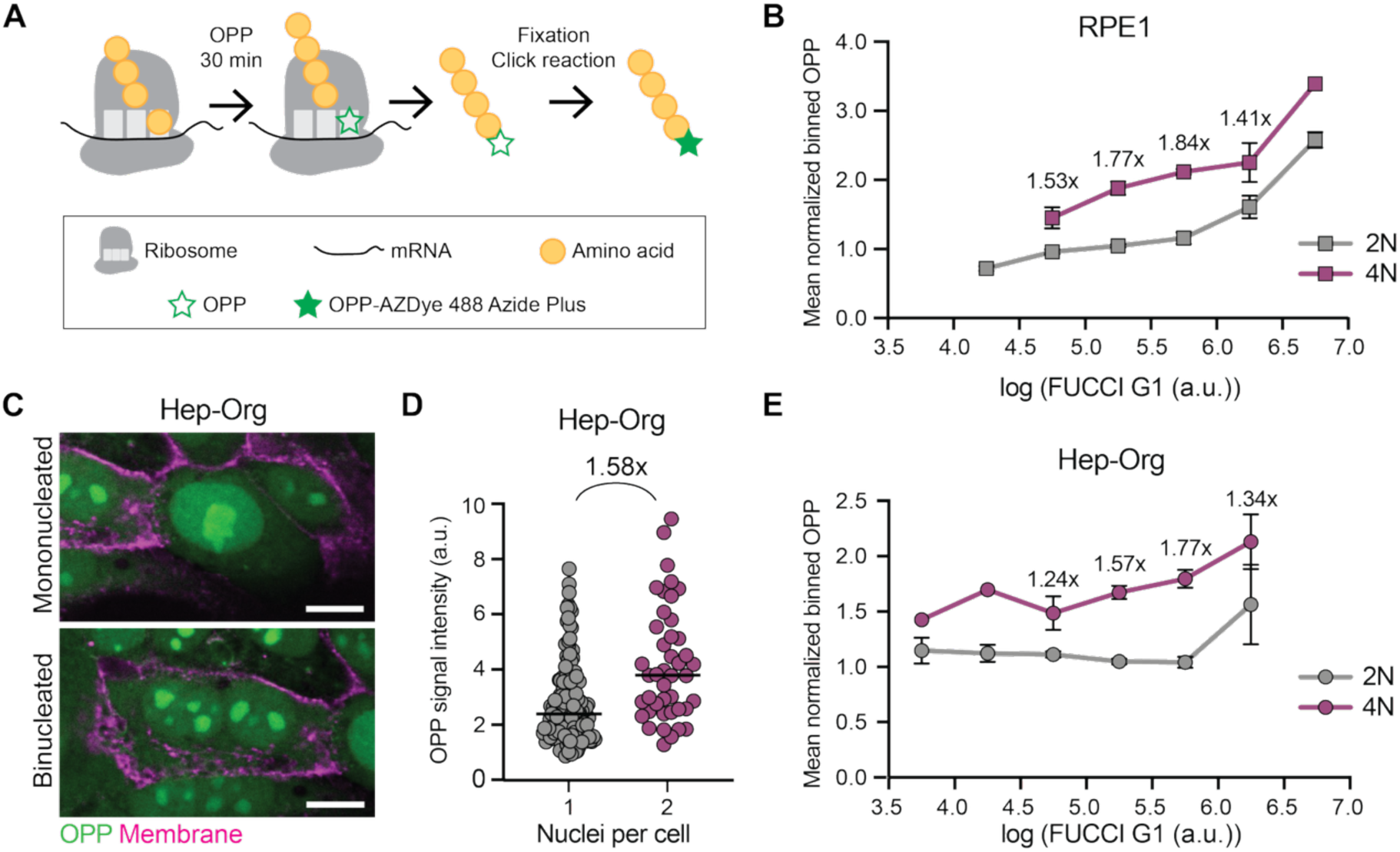
Nascent protein synthesis increases similarly with ploidy in both naturally occurring and induced polyploid cells. **(A)** Schematic of O-propargyl-puromycin (OPP) incorporation into nascent polypeptides and subsequent conjugation to a fluorescent dye by click chemistry. **(B)** Normalized OPP incorporation across G1 in diploid and induced polyploid RPE1 cells. OPP intensity was measured by calibrated flow cytometry and cells in G1 were divided into six equal bins based on log (FUCCI G1) signal. Mean normalized OPP signal is shown for 2N (gray, n > 165000 cells per replicate) and 4N (purple, n > 6000 cells per replicate) cells from N = 3 independent experiments. Error bars represent SD. **(C)** Representative images of OPP mononucleated and binucleated cells in Hep-Orgs expressing E-cadherin-tdTomato (magenta) with OPP signal (green). Images show one plane in the middle of a Hep-Org grown in a monolayer. Scale bars represent 10 µm. **(D)** Quantification of OPP signal intensity in mononucleated and binucleated cells. Each point represents one cell (n = 159 mononucleated cells and n = 43 binucleated cells from N = 3 experiments). Black lines indicate median values and fold change between median values is indicated. **(E)** Normalized OPP incorporation across G1 in Hep-Org cells. OPP intensity was measured by calibrated flow cytometry and cells in G1 were divided into six equal bins based on log (FUCCI G1) signal. Mean normalized OPP signal is shown for 2N (gray, n > 2600 cells per replicate) and 4N (purple, n > 310 cells per replicate) cells from N = 3 independent experiments. Error bars represent SD.

To determine whether the sublinear scaling of protein synthesis is a general consequence of polyploidization or specific to cells that are forced to become polyploid through cytokinesis failure, we next measured protein synthesis rates in Hep-Orgs. We reasoned that naturally occurring polyploid cells such as hepatocytes, which are developmentally programmed to become polyploid, might maintain proportional biosynthetic capacity. We first quantified OPP incorporation by imaging in Hep-Orgs expressing an endogenously-tagged E-cadherin-tdTomato membrane marker and grown as monolayers to avoid signal overlap between cells in three-dimensional structures and to ensure comparable focal plane positioning across cells, thereby enabling a quantitative comparison of OPP signals from different cells. OPP was added to the growth medium for 30 minutes prior to fixation and click-chemistry based fluorescent labeling. As previously reported, OPP signal appeared as diffuse cytoplasmic labeling with bright nucleolar foci (**Figure 3C**)^39,40^. Although total OPP signal was higher in binucleated cells compared to mononucleated cells, the median increase was only 1.58-fold (**Figure 3D**), markedly lower than the two-fold increase expected if protein synthesis scaled proportionally with ploidy. To rule out that this apparent scaling defect reflects differences in cell-cycle distribution rather than biosynthetic capacity, we repeated the OPP assay by flow cytometry in Hep-Orgs expressing the FUCCI reporter, enabling direct comparison of G1 cells across ploidy levels. Protein synthesis in naturally occurring polyploid G1 cells was increased by 1.52 ± 0.09-fold relative to diploid G1 cells (mean ± SEM, N = 3, **Figure 3E**), thus again scaling sublinearly with ploidy. Together, these results demonstrate that while polyploid cells produce more protein than diploid cells, both naturally occurring and induced polyploid cells fail to scale protein synthesis proportionally with ploidy. This suggests that after genome doubling, the scaling of protein synthesis is not inherently different between induced and naturally occurring polyploid cells.

### Naturally occurring polyploid hepatocytes compensate reduced protein synthesis by lengthening G1

Our findings so far indicate that cell size scales proportionally with ploidy in both naturally occurring and induced polyploid cells, and that protein synthesis scales sublinearly in both contexts. This raises a key question: if the immediate biosynthetic consequences of polyploidization are similar, why does polyploidy impair fitness only in cell types not normally programmed to become polyploid? One possibility is that naturally occurring polyploid cells have evolved mechanisms to compensate for initial imbalances in protein synthesis, and are nevertheless able to accumulate sufficient proteins to support replication of their increased genome. Rather than increasing the rate of protein synthesis, cells could accumulate sufficient proteins by extending the duration of G1 phase, thereby allowing more time for growth before entry into S phase. Although polyploid cells have been suggested to cycle slower^33,41^, G1 duration in naturally occurring polyploid hepatocytes has not been directly measured. We therefore quantified G1 duration in mononucleated and binucleated hepatocytes by live imaging of FUCCI markers in Hep-Orgs, defining G1 onset as the first frame after nuclear envelope breakdown (NEB), marked by rising Cdt1 signal, and G1 exit as the onset of Geminin signal at S-phase entry (**Figure 4A**). Strikingly, binucleated hepatocytes spent significantly longer in G1 than mononucleated hepatocytes (**Figure 4B**), revealing a substantial extension of G1 in naturally occurring polyploid hepatocytes.

**Figure 4.**
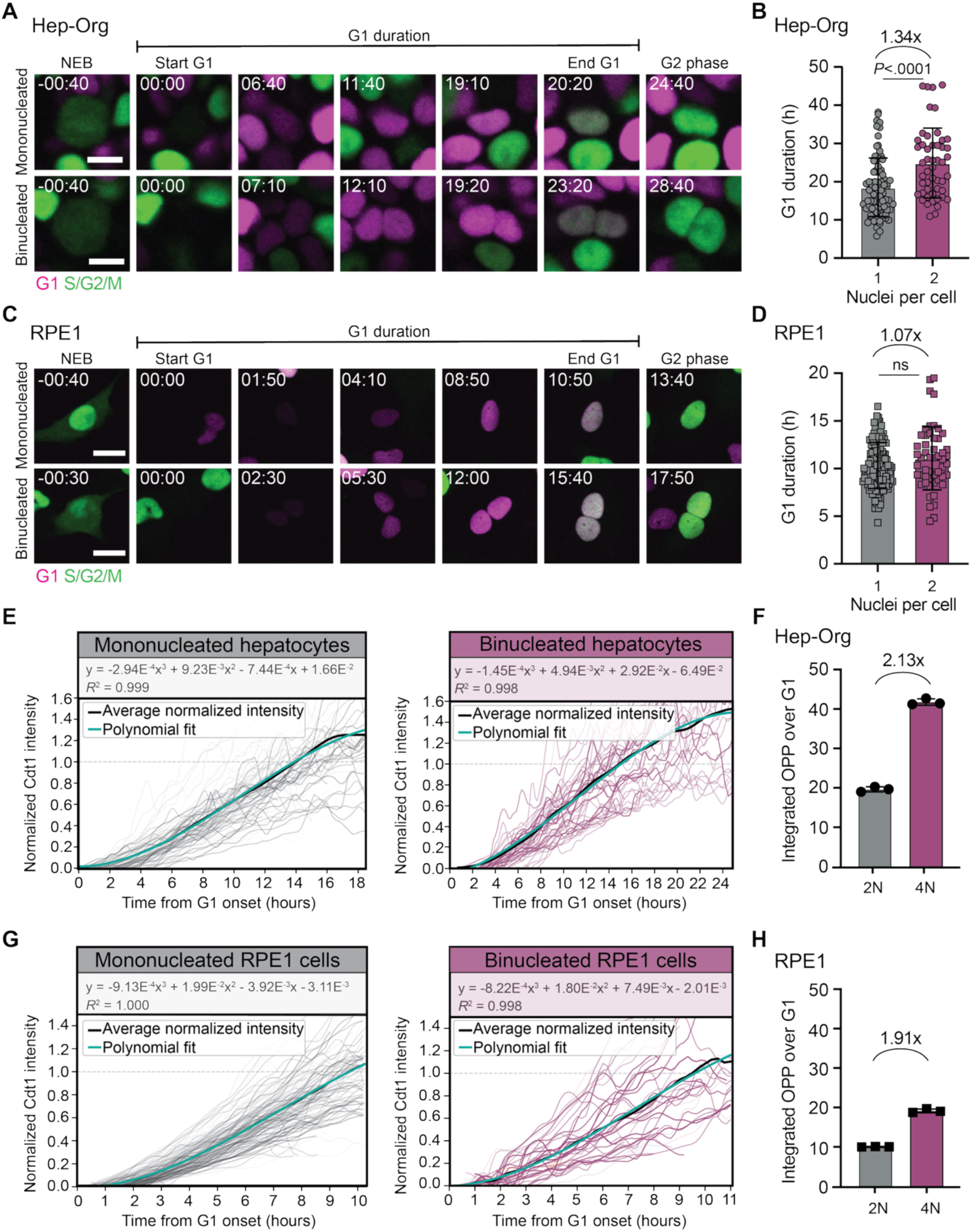
Naturally occurring polyploid hepatocytes compensate for reduced protein synthesis by lengthening G1. **(A)** Representative images of FUCCI-expressing Hep-Org cells progressing through the cell cycle. Examples of a mononucleated cell (top) and binucleated cell (bottom) are shown. Cells were imaged every 10 minutes. The start of G1 was defined as the first frame following nuclear envelope breakdown (NEB) in which Geminin (green) signal was absent. Cdt1 (magenta) progressively accumulates during G1 until Geminin rises again, marking entry into S phase. Time is indicated relative to start of G1 phase in hours:minutes. Scale bars represent 10 µm. **(B)** G1 duration in mononucleated (grey) and binucleated (purple) Hep-Org cells. Each point represents one nucleus (n = 81 mononucleated nuclei and n = 50 binucleated nuclei from N = 4 independent experiments). G1 duration is plotted in h = hours. Bars indicate mean ± SD. *P* value was calculated using a Mann-Whitney test. **(C)** Representative images of FUCCI-expressing RPE1 cells progressing through the cell cycle. Cdt1 (magenta) accumulates during G1, while Geminin (green) is absent and appears upon entry into S phase. Time is indicated relative to start of G1 phase in hours:minutes. Scale bars represent 20 µm. **(D)** G1 duration in diploid and induced polyploid RPE1 cells. Each point represents one nucleus (n = 130 mononucleated nuclei and n = 52 binucleated nuclei from N = 3 independent experiments). Bars indicate mean ± SD. ns = not significant, Mann-Whitney test. **(E)** Dynamics of Cdt1 accumulation during G1 in Hep-Orgs measured by live imaging. Normalized single-nucleus Cdt1 trajectories were aligned to the first frame after NEB. Graph on the left shows tracks of nuclei of mononucleated cells (n = 81, grey lines); graph on the right shows tracks of nuclei of binucleated cells (n = 50, purple lines). Average traces (black) were fitted using a third-degree polynomial (green). Each graph depicts data from N = 4 independent live-imaging experiments, using 10-minute intervals. **(F)** Estimated total protein production during G1 in diploid (2N, gray) and polyploid (4N, purple) Hep-Orgs. Mean normalized OPP signals (data from Figure 3E) were integrated over G1 using the fitted Cdt1-based timing calculated from live-imaging experiments. Dots indicate mean of N = 3 independent experiments and error bars indicate SD. **(G)** Dynamics of Cdt1 accumulation during G1 in RPE1 cells measured by live imaging. Normalized single-nucleus trajectories were aligned to the first frame after NEB. Left graph shows tracks from mononucleated cells (n = 130, grey lines) and right graph from binucleated cells (n = 52, purple lines) from N = 3 independent experiments, with average traces (black) were fitted using a third-degree polynomial (green). **(H)** Estimated total protein production during G1 in diploid (2N, gray) and induced polyploid (4N, purple) RPE1 cells, calculated as in Panel F, using data from Figure 3B. Dots indicate mean of N = 3 independent experiments and error bars indicate SD.

To determine whether induced polyploid cells undergo a similar adaptation, we measured G1 duration in induced tetraploid RPE1 cells using live imaging of the FUCCI reporter (**Figure 4C**). In contrast to hepatocytes, induced polyploid RPE1 cells showed no significant increase in G1 duration among those that progressed into S phase (**Figure 4D**), consistent with previous reports^24^. Notably, 42% of induced binucleated cells (N = 3) failed to enter S phase within 20 hours of imaging, indicating G1 arrest rather than delayed progression. Together, these results reveal a striking divergence: naturally occurring polyploid hepatocytes substantially extend G1, whereas induced polyploid RPE1 cells either arrest in G1 or progress into S phase without extending G1.

We next asked whether the G1 extension observed in polyploid hepatocytes is sufficient to compensate for their sublinear protein-synthesis scaling and to restore total protein production to levels proportional to their increased ploidy. To estimate cumulative protein production during G1, we first established a relationship between Cdt1 levels and relative G1 progression by fitting a polynomial to average Cdt1 accumulation trajectories obtained from live imaging of single mononucleated and binucleated hepatocytes (**Figure 4E, S4A**), as described previously^35^. This model allowed us to use Cdt1 levels measured by flow cytometry as a proxy for relative G1 time, enabling us to assign OPP-based translation rates to specific timepoints within G1. Integrating these translation rates over the full duration of G1 then allowed us to estimate the total amount of protein produced over the course of G1 in diploid and polyploid hepatocytes (**Figure 4F**). This analysis revealed that the combined effect of increased translation rates and extended G1 duration is sufficient to restore total protein production in polyploid hepatocytes to levels that scale proportionally with ploidy (2.13-fold, **Figure 4F**). Moreover, G1 lengthening is estimated to account for approximately 39% of this increase, highlighting it as a major contributor to biosynthetic compensation (**Figure S4B).** Applying the same analysis to RPE1 cells revealed that despite the modest increase in translation rates in induced polyploid cells, the minimal G1 extension in these cells resulted in lower total protein production during G1 compared to naturally occurring polyploid hepatocytes (1.91-fold, **Figure 4G, H and S4B**). Thus, unlike hepatocytes, induced polyploid RPE1 cells fail to sufficiently extend G1 to compensate for reduced protein synthesis scaling, leaving them with a cumulative biosynthetic deficit relative to their increased genome content.

### The hepatocyte response to WGD is the same whether polyploidy arises naturally or is induced by cytokinesis failure

The limited G1 extension observed in induced polyploid RPE1 cells contrasts with the robust G1 lengthening seen in naturally occurring polyploid hepatocytes, suggesting that these divergent responses reflect either the mode of polyploidization (endomitosis versus induced cytokinesis failure) or intrinsic properties of the cell type itself. To distinguish between these possibilities, we experimentally induced polyploidy in Hep-Orgs by a short DCB treatment and compared cell size and protein synthesis efficiency between induced and naturally occurring polyploid hepatocytes using calibrated flow cytometry (**Figure 5A**). Although induced and naturally occurring polyploid cells cannot be distinguished within the population, DCB treatment reproducibly increased the percentage of 4N cells relative to DMSO-treated control cells (**Figure 5B**). Despite a small increase in diploid cell sizes across G1 after DCB treatment, DCB-treated polyploid hepatocytes exhibited cell volumes comparable to those of naturally occurring polyploid cells (**Figure 5C**). Similarly, there is no clear difference in protein synthesis efficiency between naturally occurring and induced 4N hepatocytes (**Figure 5D**).

**Figure 5.**
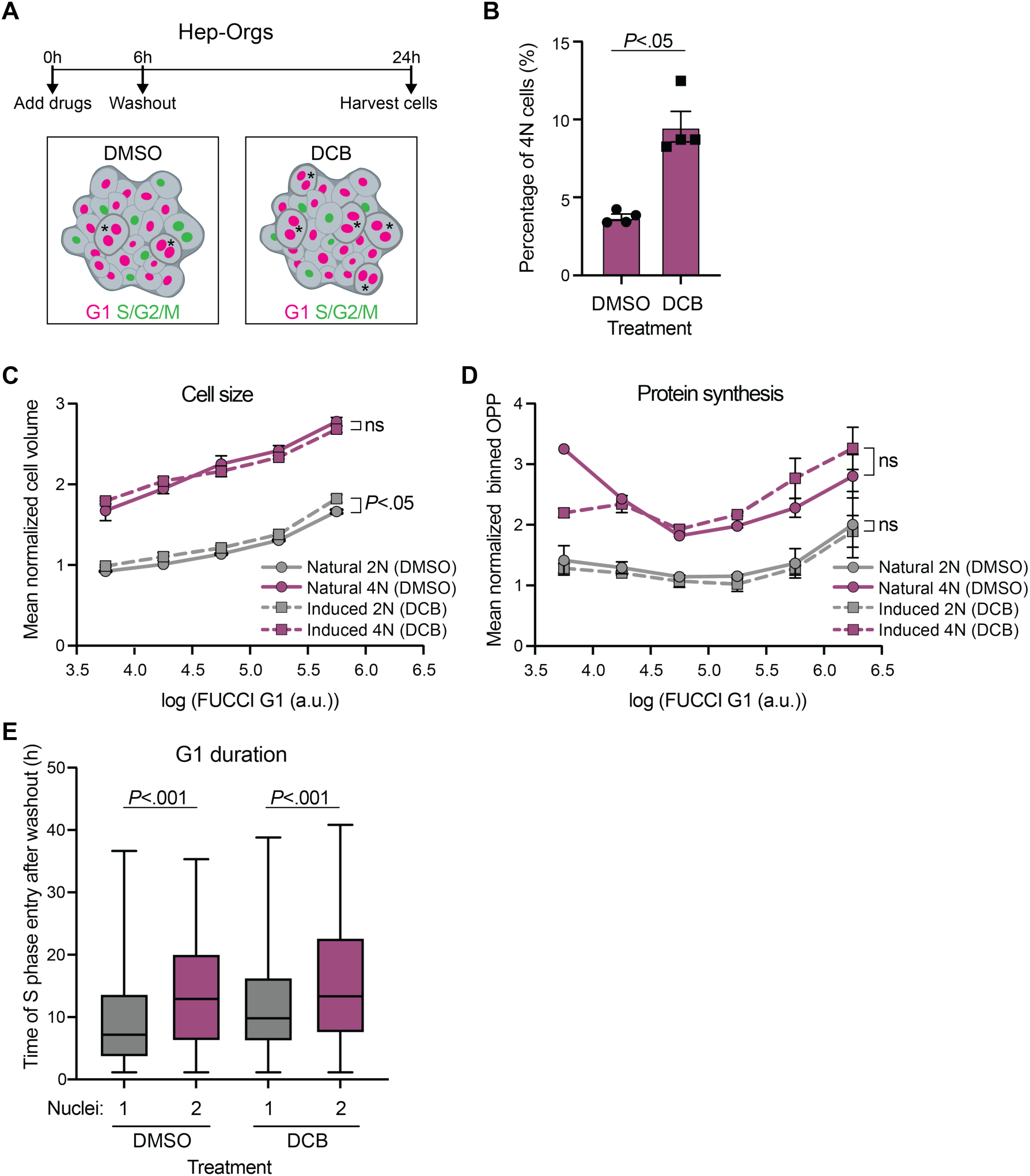
Hepatocytes scale cell size, protein synthesis, and G1 duration with ploidy independently of how polyploidy arises. **(A)** Schematic of experimental workflow to induce polyploidy in Hep-Orgs. Cells were treated with DMSO or DCB for 6 hours, after which the drug was washed out, and the newly formed polyploid cells were allowed to progress through G1 phase for 18 hours before harvesting cells (h = hours, asterisks mark binucleated cells in G1). **(B)** Percentage of 4N cells in Hep-Orgs measured by flow cytometry following treatment with DMSO or DCB. Bars indicate mean ± SEM (N = 4 independent experiments, n > 3500 per sample). *P* value was calculated using a paired Student’s *t*-test. **(C)** Normalized cell volumes measured by flow cytometry of naturally occurring (DMSO-treated) and induced (DCB-treated, dashed lines) 2N and 4N hepatocytes. Cells in G1 were binned based on log(FUCCI G1) signal and mean normalized cell volume is shown for 2N (gray) and 4N (purple) cells. N = 3 independent experiments. Error bars represent SEM. Statistical comparisons were performed using paired Student’s *t*-tests; ns, not significant. **(D)** Nascent protein synthesis scaling with ploidy in naturally occurring and induced Hep-Orgs. Mean normalized OPP signal is shown for 2N (gray) and 4N (purple) cells across G1 bins from N = 2 independent experiments. Error bars represent SEM; statistical comparisons were performed by paired two-tailed Student’s *t*-tests (ns, not significant). **(E)** Time of S-phase entry in mononucleated (gray) and binucleated (purple) Hep-Org cells following DMSO or DCB treatment, measured by live imaging. Data are from N = 2 independent experiments. Boxes indicate the interquartile range (25th-75th percentile), and the central line indicates the median. Whiskers extend to the minimum and maximum values. Statistical comparisons were performed using Mann-Whitney tests.

Finally, we investigated whether the G1 lengthening observed in naturally occurring polyploid hepatocytes depends on how polyploidy arises. To address this, we compared G1 duration between naturally occurring and induced polyploid hepatocytes by live imaging of FUCCI markers following DCB washout. Strikingly, induced polyploid hepatocytes extend G1 to a similar degree as naturally occurring binucleated cells, with both groups showing significantly longer G1 duration compared to their mononucleated counterparts (**Figure 5E, S5A**). Although S-phase entry is modestly delayed in both mononucleated and binucleated cells following DCB treatment, the relative extension of G1 in binucleated hepatocytes is preserved regardless of treatment condition (**Figure 5E, S5B-D**). Together, these findings show that G1 extension following polyploidization is a hepatocyte-intrinsic response, independent of the mechanism by which polyploidy is established. The inability of RPE1 cells to similarly extend G1 therefore reflects cell-type-specific wiring rather than the mode of polyploidization.

## Discussion

Polyploid cells are widespread across development and disease, yet it remains unclear whether they share a common cellular response to WGD or whether outcomes depend on how, and in which cell type, polyploidy arises. In this study, we used three model systems, *C. elegans* intestinal cells, human hepatocyte organoids, and induced polyploid RPE1 cells, to directly compare the immediate consequences of polyploidization on cell size, protein synthesis, and cell-cycle progression. Across all systems, cell size scales proportionally with ploidy, and nascent protein synthesis scales sublinearly. Despite these similar immediate responses, naturally occurring polyploid hepatocytes compensate for their reduced protein synthesis rates by substantially lengthening G1, while induced polyploid RPE1 cells fail to do so. Importantly, when polyploidy is induced in hepatocytes by transient DCB treatment, cells still extend G1 to the same extent as their naturally occurring polyploid counterparts. Together, these findings suggest that the immediate cellular consequences of WGD are largely conserved, and that cell-type-specific cell-cycle adaptations, rather than the mode of polyploidization, determine whether polyploidy is tolerated.

Cell size and DNA content are tightly coupled, and how they scale relative to each other is an important determinant of cellular function^42^. For example, the nuclear-to-cytoplasm ratio is a key determinant of cell-cycle speed and checkpoint strength, and increases in cell size without accompanying DNA increases have been shown to promote senescence^42–48^. Although cell size has been shown to increase with increasing ploidies in many naturally occurring polyploid cells (reviewed in ^1^), a systematic analysis of how cell size and ploidy scales within single polyploid cells has been largely lacking. Here, we measured cell volume of tetraploid hepatocytes and induced tetraploid RPE1 cells in the first cell cycle after they become polyploid. Strikingly, we find that both types of polyploid cells have very similar growth rates during G1, that do not differ substantially from their diploid counterparts: both naturally and induced tetraploid cells are on average twice as big as diploid cells when they are born, and they grow similarly throughout G1. Thus, our results suggest that polyploid cells, irrespective of how they arise, do not show inherent differences in how they grow compared to diploid cells.

Interestingly, a recent study showed that cell and nuclear volume do not always scale with DNA content after WGD in cancer cells, generating populations of small and large 4N cells with distinct mitotic and tumorigenic properties^49^. In these experiments, stable 4N cancer cell clones were generated by treating cells with cytokinesis inhibitors, and selecting surviving 4N clones after multiple cell cycles. The finding that in these 4N cancer cell clones distinct populations of cell sizes can arise, suggest that despite that WGD may not have immediate consequences for cell growth, the extent by which polyploid cells grow over longer periods of time will likely vary across cell types and conditions, influencing their fitness, mitotic fidelity, and tumorigenic potential.

One important determinant of cellular fitness is the capacity to scale protein synthesis in growing cells. How protein synthesis scales with ploidy has remained an open question, with some contradictory observations across systems. In budding yeast, mRNA and protein abundance scale sublinearly with ploidy, with tetraploid cells producing only ∼3-fold rather than 4-fold more protein than haploids, due to reduced ribosomal biogenesis and protein translation^23^. In contrast, lymphocytic leukemia cells maintain a constant mass-normalized growth rate across a 100-fold range of polyploidization^50^, and our unpublished observations in *C. elegans* intestinal cells indicate that protein translation becomes increasingly efficient at higher ploidies (above 16C, C. Jordan Ortiz and M. Galli, unpublished data). Again, these divergent responses of polyploidization on biosynthesis could reflect differences in the mode of polyploidization, the cell type, or the extent to which cells have had time to adapt to their increased ploidy. Our finding that nascent protein synthesis scales sublinearly with ploidy in both naturally occurring and induced mammalian polyploid cells indicates that, at least in the immediate cell cycle following WGD, mammalian cells resemble yeast in their inability to fully scale biosynthesis with ploidy.

Although protein synthesis is not proportionally scaled to ploidy immediately after WGD, naturally occurring polyploid hepatocytes appear to accommodate this imbalance by extending G1, providing additional time to accumulate biomass before DNA replication. Because most cell growth occurs in G1, lengthening this phase allows increased biosynthesis without the need to alter protein synthesis rates. How exactly DNA content is sensed by the cell-cycle machinery to extend G1 duration after WGD, and why this does not always occur after induction of cytokinesis failure in RPE1 cells is not completely understood. A p53-dependent tetraploidy checkpoint has long been recognized in non-transformed cells, in which cytokinesis failure leads to elevated p21 levels and stable G1 arrest^51–54^. Increased DNA content is sensed indirectly due to the presence of extra centrosomes in cells that have undergone a WGD, and these additional centrosomes activate downstream signaling through the Hippo pathway or the PIDDosome to stabilize p53 and induce a p21-mediated cell-cycle arrest^53,54^. Cell-to-cell variability in p53 activity is thought to determine whether individual tetraploid cells arrest or progress through the cell cycle^53–57^. Our observation that 42% of induced binucleated RPE1 cells fail to enter S phase, while the remainder progress without extending G1, could thus be explained by heterogeneity in p53 stabilization, allowing many cells to bypass arrest despite their polyploid state. It is possible that polyploid hepatocytes, which are developmentally programmed to become polyploid, possess intrinsic mechanisms that couple ploidy to G1 length, which are not present in RPE1 cells, leading to cells being either stuck in G1 or entering S phase without a G1 extension. Identifying the molecular basis of this difference will be essential for understanding why some cell types tolerate polyploidy while others do not.

Several limitations should be considered when interpreting our results. First, we only focused our analyses on *C. elegans* intestinal cells and two mammalian cell types, hepatocytes and RPE1 cells, and how broadly our findings generalize to other polyploid contexts remains to be determined. Mammalian hepatocytes are one of the few naturally polyploid cell types that retain proliferative capacity in vivo, allowing them to divide after becoming polyploid. Many naturally occurring polyploid cells, including *C. elegans* intestinal cells, continue endoreplication cycles, but do not enter M phase after becoming polyploid. Whether cell growth is different in polyploid cell types that continue to proliferate versus cells that terminally differentiate remains to be determined. Second, our analyses of polyploid cell growth and protein synthesis were restricted to the first cell cycle following polyploidization, and we did not examine long-term consequences after multiple divisions. Nonetheless, replication stress and genetic instability have been shown to arise already in the first S phase after induced WGD^24^, supporting the idea that the immediate cellular response shapes longer-term outcomes. To our knowledge, this work is the first to directly compare induced and naturally occurring polyploid cells side by side, linking cell size, protein synthesis, and cell-cycle progression at the single-cell level immediately after WGD.

Our findings have implications for both physiological and pathological polyploidy. In developmental contexts, where polyploidy is associated with specialized functions such as enhanced biosynthesis and tissue growth, the ability to compensate for reduced protein synthesis through G1 lengthening may be essential to ensure that polyploid cells accumulate sufficient biomass to support their increased genome and their physiological roles. In contrast, induced polyploid cells fail to initiate this response, leaving them with a cumulative biosynthetic deficit and an uncoupling of growth from cell-cycle progression, which has been shown to underlie their reduced fitness by inducing chromosomal instability in the first S phase after WGD^24^. These differences are particularly relevant for cancer, where WGD is a frequent early event and is associated with chromosomal instability, metastasis, and poor prognosis^58–61^. Our work suggests that whether a polyploid state is tolerated depends less on the mode of polyploidization and more on whether cells can lengthen their cell cycle to match their increased genome.

## Materials and methods

### C. elegans strains and maintenance

Strains were maintained at 15°C and grown on agar NGM (Nematode Growth Medium) plates seeded with OP50 *Escherichia coli* bacteria according to standard protocols. Strains used for this study are listed in **Table S1**. The intestine and body-wall muscle specific membrane (*matIs159*) transgenes were generated by gamma irradiation of an extrachromosomal array containing a plasmid with *ges-1p::GFP-PH* and *myo-3p::mCherry-PH* sequences together with a *lin-48p::TdTomato* co-injection marker. *AID::mcm-4* (*mat106*) and *AID::mcm-7* (*mat138*) alleles were generated using CRISPR/Cas9-based genome editing as previously described^62^ and crossed to the *ges-1p::TIR1* (*ieSi61*) allele^31^ and strain with intestinal and body-wall membrane markers to generate the strain used for ploidy reduction experiments.

### RPE1 and bepatocyte organoid culturing

Cells were maintained at 37°C in a 5% CO_2_ atmosphere. RPE1 FUCCI (mkO2-hCdt1-mAG-hGeminin) cells^35^ were maintained in DMEM/Nutrient Mixture F-12 (DMEM/F12, GlutaMAX supplement, Gibco, #31331) supplemented with 10% fetal bovine serum (FBS, Sigma-Aldrich, #F7524) and 1% penicillin/streptomycin (Gibco, #15140). Hep-Orgs used in this study were previously generated from human fetal livers with no objection from the Ethical Medical Council of the Leiden University Medical Center (LUMC)^26^. The Hep-Org line expressing endogenous E-cad-herin-tdTomato was generated using CRISPaint as previously described^27,28,63^. All organoids were grown in Cultrex reduced growth factor basement membrane extract (BME), type 2 (R&D Systems, #3533-001) and cultured and passaged as previously described^26^. Culture medium components are listed in **Table S2**. Culturing medium was refreshed every 2 or 3 days, and organoids were passaged every 7-10 days at a 1:3-1:6 ratio by manual pipetting with a P1000 pipette. During the first few days of organoid line establishment the culturing medium was supplemented with extra Y-27632 (final concentration: 10 μM) to minimize anoikis. All cell lines used in this study were regularly tested for mycoplasma and were negative.

### Generation of Hep-Org FUCCI line

For the generation of the hepatocyte organoid line expressing the FUCCI reporter, the FUCCI construct (Clover-Geminin-IRES-mKO2-Cdt1-WPRE, Addgene #83841) was amplified and cloned into a the piggyBac donor plasmid between terminal repeats using Gibson assembly. Correct insertion was confirmed by Sanger sequencing. Human hepatocyte organoids were transfected by electroporation using a two-plasmid piggyBac transposon system consisting of a transposase plasmid and the donor plasmid. Electroporation was performed as described previously^64^, with the addition of 1.25% DMSO (Sigma-Aldrich, #D8418) to the cells 24 hours prior to electroporation^65^. After electroporation, outgrowing fluorescent organoids were manually picked, dissociated into single cells with Accutase™ (Gibco, #A1110501), and plated into a single BME droplet per well of a 48-well plate for clonal expansion. At least two independent clonal FUCCI lines were generated and used for downstream analyses.

### C. elegans intestinal cell ploidy reduction and quantification

For ploidy reduction experiments, NGM plates were supplemented with 0.5 mM auxin (Sigma-Aldrich, #I2886) and allowed to dry at room temperature for three days. Control plates without auxin, were supplemented with equal volumes of 100% ethanol. To obtain synchronized populations of L1 larvae, NGM plates containing gravid adults were first washed in M9 buffer. After washing, the worms were bleached using a buffer containing 0.5 M NaOH and 5% NaClO for 5-7 minutes. The eggs were then washed and left in M9 buffer overnight at room temperature to allow the eggs to hatch. The isolated embryos were then plated and grown on NGM plates with and without auxin (+ and ct). Synchronized worms were grown at 15°C for 56 hours corresponding to L4 stage. Worms were subsequently washed off with Milli-Q and placed in slides coated with poly-L-lysine (Sigma-Aldrich, #P8920). Worm cuticles were removed by freeze-cracking and placed in Carnoy’s fixation solution (60% EtOH, 30% acetic acid, 10% chloroform) for one hour. The worms were then treated with 100μL of 20 μg/mL RNAse A solution (Sigma-Aldrich, #10109142001) and incubated for one hour at 37°C. This was followed by staining with 100 μg/mL of propidium iodine (PI) (Sigma-Aldrich, #P4170) and incubated at room temperature for 60 minutes in a humidified chamber. Slides were then washed three times and mounted with ProLong Gold Antifade (Invitrogen, #P36934) mounting agent. Images were captured with a Nikon Ti2 microscope equipped with a CSU-X1 spinning disk scanner unit and a CMOS digital camera (C13440; Hamamatsu Photonics, Hamamatsu-city, Japan). Intestinal cellular ploidy levels were quantified using FIJI software by measuring the integrated density of the PI signal of the nuclei of the third intestinal ring. The signal was then normalized to the average signal of three proximal body-wall muscle (2C) nuclei.

### Cell volume measurements of C. elegans larvae

To measure cell volumes, worms were bleach-synchronized as described above, after which they were plated on NGM plates and grown at 15°C degrees. Worms were harvested at the beginning of each larval stage (after 2 hours for L1 stage, 29 hours for L2 stage, 43 hours for L3 stage and 56 hours for L4 stage) for measurements of cell volumes in wildtype animals (using strain GAL316). For measurements of cell volume after ploidy reduction, synchronized GAL232 L1 larvae were grown on either control or auxin NGM plates at 15°C degrees for 56 hours (L4 stage). For imaging, worms were mounted on 4% agarose pads and immobilized with 1 or 2 mg/mL of tetramisole (Sigma Aldrich, #L9756). A 22 mm coverslip was overlaid and sealed with VALAP (a 1:1:1 mixture of vaseline, lanolin and paraffin). Images were acquired on a Nikon Ti2 microscope equipped with a CSU-X1 spinning disk scanner unit and a CMOS digital camera (C13440; Hamamatsu Photonics, Hamamatsu-city, Japan) with 100x oil objective for L1 and L2 larval stages and at 60x oil objective for L3 and L4 larval stages. Confocal image stacks were processed and analyzed using Imaris (version 10.0.1; Bitplane, Zurich, Switzerland). Cells were segmented using the Surfaces module, and volume measurements were extracted for quantitative analysis.

### Induction of polyploid cells in RPE1 cells and Hep-Orgs

To induce polyploid cells, RPE1 and Hep-Orgs expressing the FUCCI markers were incubated with either 4 µM (RPE1 cells) or 5 µM (Hep-Orgs) Dihydrocytochalasin B (DCB, Cayman Chemical, #CAYM20845) in the culture medium. Control cells were treated for the same duration with the equivalent concentration of DMSO (Sigma-Aldrich, #D8418) in the corresponding culture medium. Following treatment, cells were washed three times with PBS and then supplied with fresh culture medium. The duration of drug treatment and recovery depended on the experiment and is specified in the results section and figures.

### Cell size measurements Hep-Orgs by imaging

Cell size measurements were performed in Hep-Orgs expressing endogenous E-cadherin-tdTomato^27^ and stained with DRAQ5 (1:1,500, BioLegend, #424101). Imaging was performed on live organoids using a Nikon TiE microscope equipped with Borealis illumination unit (Andor), CSU-W1 spinning disk scanner (Yokogawa), and iXon-888 Ultra EMCCD camera (Andor) using a UPLSAPO-S 30x silicone objective (Olympus) with 1.5x optical zoom in an environmentally-controlled chamber held at 37°C with 5% CO_2_. Whole organoids were imaged with 0.5 μm z-intervals. Cell volumes were calculated using a prismatoid model based on the cross-sectional areas of the top, middle, and bottom planes (**Figure S1D**). Area measurements were taken using FIJI^66^.

### Cell size-calibrated flow cytometry

To estimate cell diameters for volume calculations, a calibration curve of the forward scatter (FSC) was generated using a CytoFLEX (*Beckman Coulter)*. Polymer microspheres with defined diameters (6, 12, 17, 19 and 23 µm, Thermo Fisher, #7505A), spanning the size range of the measured cells, were used to obtain reference FSC-A values. The calibration curve was generated from three replicate measurements collected over the period during which the experiments were performed (**Figure S1E**). This curve was then used to estimate cell volumes from FSC-A measurements obtained by flow cytometry. Because scattering properties vary between cell types, this approach is only suitable for comparisons between populations within the same sample, in this case diploid and polyploid cells.

For cell volume measurements of induced polyploid cells, RPE1 cells were incubated with DCB for approximately 4-5 hours, after which the drug was washed out. Cells were harvested after a total of 9 hours. For cell volume measurements of induced polyploid cells in Hep-Orgs, organoids were incubated with DMSO or DCB for 6 hours, after which drugs were washed out, and cell volumes were measured 24 hours after induction.

RPE1 cells and Hep-Orgs were dissociated into single cells using TrypLE (Gibco, #12605) according to standard protocols. Cells were washed with PBS, filtered, and stained with Hoechst 34580 (Sigma-Aldrich, #63493) at a final concentration of 1 µg/mL for RPE1 cells and 10 µg/mL for Hep-Orgs. After incubation for 15 minutes at 37°C and 5% CO_2_, cells were analyzed on a CytoFLEX (Beckman Coulter). G1 cells were gated based on the FUCCI marker and Hoechst signal was used to distinguish 2N and 4N populations. Gated populations were exported with FlowJo™ v10.10 Software (BD Life Sciences), and cell volumes were calculated from FSC.A values using the calibration curve and further analyzed by custom Python (v3.12.7) scripts. Calculated cell volumes were normalized per replicate to the average calculated volume of 2N cells in mid-G1.

### Nascent protein synthesis assay in Hep-Orgs by imaging

To measure protein synthesis rates, Hep-Orgs were pulse-labeled with the puromycin analog O-Propargyl-puromycin (OPP), which incorporates into nascent polypeptide chains. Hep-Orgs were dissociated into single cells using TrypLE (Gibco, #12605) and plated onto ibidiTreat 8-well chambered coverslips (ibidi, #80826) overnight to allow attachment as a monolayer. OPP assay was performed using Click-&-Go Plus 488 OPP Protein Synthesis Assay Kit (Click Chemistry Tools, #1493). In short, cells were labeled with 20 µM OPP in culture medium for 30 minutes, fixed in 4% paraformaldehyde in PBS (Electron Microscopy Sciences, #RT15710) for 10 minutes, and permeabilized with 0.5% Triton-X100 (Sigma-Aldrich, #T8787) in PBS for 20 minutes. The click reaction was performed according to the manufacturer’s protocol, after which cells were washed thoroughly with PBS and co-stained with DAPI (0.25 ng/mL, Sigma-Aldrich, #32670). Cells were rinsed and imaged in PBS using a Plan Apo λ 60x oil objective (Nikon, #MRD01605) on a Nikon Ti2 microscope equipped with an L6Cc illumination unit (Oxxius), CSU-X1 spinning disk scanner unit (Yokogawa), and with a C11440-22C camera (Hamamatsu). Whole organoids were imaged with 0.5 µm z-intervals. OPP signal was quantified from sum projections of corresponding z-slices by measuring the integrated density using FIJI^66^ in individual cells.

### Nascent protein synthesis assay by flow cytometry in RPE1 cells and Hep-Orgs

To perform the OPP assay on induced polyploid cells, RPE1 FUCCI cells were treated with DCB for approximately 2,5 hours, after which the drug was washed out. Cells were then incubated with OPP and fixed 6 hours after the start of induction. To analyze OPP in induced polyploid cells in Hep-Orgs, cells were treated with DMSO or DCB for 6 hours, followed by washout and incubation in fresh culture medium. After OPP incubation, cells were fixed approximately 25 hours after the start of induction. The OPP assay was performed similarly to the imaging protocol described above, except that all steps were carried out on single-cell suspensions in tubes. As the assay was performed in cells expressing FUCCI markers, OPP was labeled using the far-red dye AZDye647-Picolyl-Azide (Jena bioscience, #CLK-1300A-AZ) according to manufacturer’s protocol. Cells were washed, resuspended in PBS, and analyzed on a CytoFLEX (Beckman Coulter). G1 cells were gated based on the FUCCI markers, and Hoechst signal was used to distinguish 2N and 4N populations. Gated populations were exported using FlowJo™ v10.10 Software (BD Life Sciences) and further analyzed by custom Python (v3.12.7) scripts. OPP signal was normalized per replicate to the average signal of 2N cells in mid-G1.

### Long-term live imaging of Hep-Orgs and RPE1 cells

For live imaging, RPE1 FUCCI cells or FUCCI-expressing Hep-Orgs were plated onto ibidi 8-well chambered coverslips with ibiTreat (ibidi, #80826) at least one day prior to imaging. To induce polyploid cells in RPE1 cells, DMSO or DCB was added for approximately 2.5 hours, after which the drug was washed out, and live imaging was continued at the same positions. For live imaging of induced polyploid Hep-Orgs, organoids were incubated with DMSO or DCB for approximately 5 hours on the day of the experiment, followed by washing and replacement with fresh medium. Live imaging was started approximately 1 hour later, so 6 hours after the start of DCB treatment.

Fluorescent excitation was kept to a minimum. Imaging was performed using a Nikon TiE microscope equipped with Borealis illumination unit (Andor), CSU-W1 spinning disk scanner unit (Yokogawa), and an iXon-888Ultra EMCCD camera (Andor), using a UPLSAPO-S 30x silicone objective (Olympus) in an environmentally controlled chamber held at 37°C with 5% CO_2_. RPE1 cells were imaged using NIS-elements software (Nikon) for 20-25 hours with 10-minute intervals with approximately 7 z-slices spaced 1 µm apart. Organoids were imaged for 40–65 hours with 10-minute intervals, with 12 z-slices spaced 5 µm apart per position.

### Analysis of live imaging of Hep-Orgs and RPE1 cells

Cell tracking was performed using the TrackMate plugin^67^ in FIJI. Background signal was measured in regions outside cells or organoids and subtracted from all measurements. Tracks were exported and further analyzed using custom Python (v3.12.7) scripts. Using napari^68^, the start of G1 was manually annotated as the time point immediately following nuclear envelope breakdown, when Geminin signal was absent. Tracks were also annotated as coming from mononucleated or binucleated cells. The end of G1 was determined automatically by identifying the time point of the steepest increase in geminin signal. To reduce variability, raw tracks were smoothed using a Savitzky–Golay filter (7-point window, second-order polynomial). G1 duration was calculated as the time between the annotated start of G1 and the onset of Geminin expression.

To convert raw Cdt1 signal measured by flow cytometry into time, live-imaging tracks were normalized per replicate to the Cdt1 signal of cells in early S phase. Smoothed Cdt1 signals from individual tracks were averaged separately for mononucleated and binucleated cells. A third-order polynomial was then fitted to the average Cdt1 signal for each population, using only the data up to the average G1 duration of that population. Cdt1 signals measured by flow cytometry were similarly normalized to Cdt1 signal from early S phase cells per replicate. Time was inferred independently for each population using the formula of the fit, enabling plotting of OPP signal of cells in G1 phase over time. Accumulated protein synthesis during G1 was calculated as the area under the curve from 0 hours to the average G1 length of each population.

Live imaging data of Hep-Orgs treated with DMSO or DCB used for Figure 5 were analyzed separately. Full G1 duration could not be determined because imaging did not include the treatment period. To enable a fair comparison of estimated G1 duration between mononucleated and (induced) binucleated Hep-Orgs, only cells that were already in G1 at the start of imaging were included, ensuring that all analyzed cells had been exposed to DMSO or DCB prior to imaging. Consequently, cells born during imaging were excluded from the analysis. Entry into S phase was manually annotated for all cells using the Cell Counter plugin in FIJI.

### Data analysis

All images were processed and analyzed using FIJI (Schindelin et al., 2012). Most graphs and statistical analysis were conducted in GraphPad Prism version 11.0.0. Some data processing and the graphs with live-imaging tracks were made in custom Python (v3.12.7) scripts using NumPy (v2.2.6), pandas (v2.2.3), matplotlib (v3.10.3), and seaborn (v0.13.2). A minimum of two biological replicates were conducted for each experiment. Statistical significance was determined using paired or unpaired Student’s *t*-tests, or Mann-Whitney tests, depending on data distribution.

## Acknowledgments

We thank M. Tanenbaum (Hubrecht Institute) for sharing the RPE1-FUCCI cell line, members of the Clevers lab for organoid culturing advice and reagents, A. Pfauth from the Hubrecht FACS facility for support with setting up the size calibrated flow cytometry, R. van der Palen and N. Perry Levy for generating *C. elegans* strains, and A. de Graaff and P. Toonen from the Hubrecht Imaging Centre for microscopy support. The *ieSi61* allele was provided by *Caenorhabditis* Genetics Center (CGC), which is funded by NIH Office of Research Infrastructure Programs (P40 OD010440). This work was supported by funding from the Cancer Genomics Centre (cancergenomics.nl) and an NWO-M1 grant (OCENW.KLEIN.258) from the Dutch Research Council (NWO).

