## Supplemental Figures for "The immediate cellular response to whole-genome doubling is conserved across polyploid contexts"

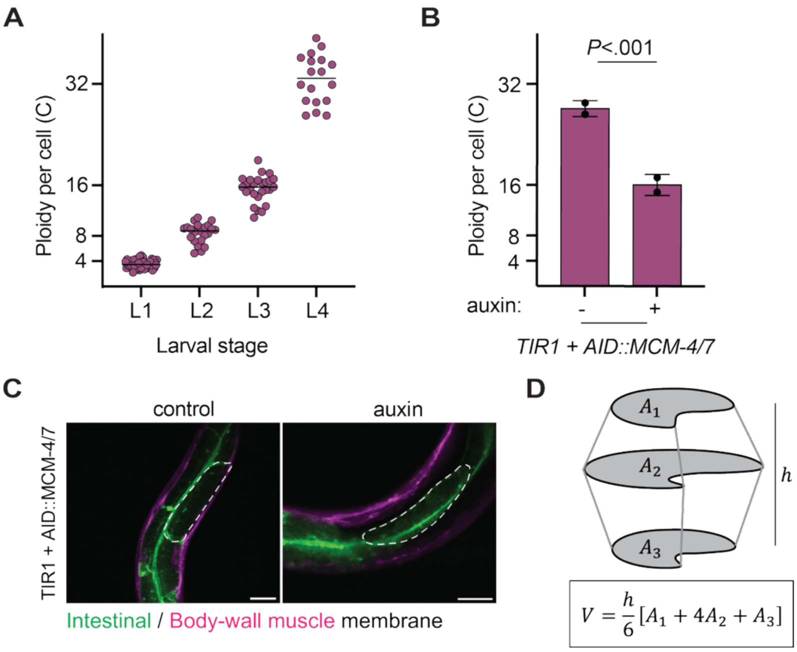


**Figure S1. Quantification of cellular ploidy levels and cell volumes.**

**(A)** Quantification of intestinal cell ploidies in *C. elegans* larval stages in animals expressing GFP-PH and mCherry-PH. Each dot represents the cellular ploidy in one animal (at least 19 worms were quantified per larval stage). Data are from N = 1 experiment. **(B)** Quantification of intestinal cellular ploidy in L4 stage for *Pges-1::Tir1, AID::mcm-4, AID::mcm-7* strain in control (ct, n = 40, left) and reduced ploidy (+ auxin, n = 39, right). Bars and error bars indicate mean ± SD. Individual points show N = 2 independent replicates. Statistical analysis was performed using a two-tailed Student's *t* test. **(C)** Representative images of live imaging of GFP-PH (intestinal membrane, green) and mCherry-PH (body-wall muscle membrane, magenta) in intestinal ring 3 for *Pges-1::Tir1, AID::MCM-4, AID::MCM-7* strain in control (ct, n = 30) and reduced ploidy (+ auxin, n = 30) used for quantifications in **Figure 1D**. Dashed lines indicate cell outlines. Scale bar is 20 µm. **(D)** Schematic of cell volume estimation in Hep-Orgs using a prismatoid using area measurements of top, middle, and bottom plane of a cell and the formula to calculate cell volume plotted in **Figure 1G**.

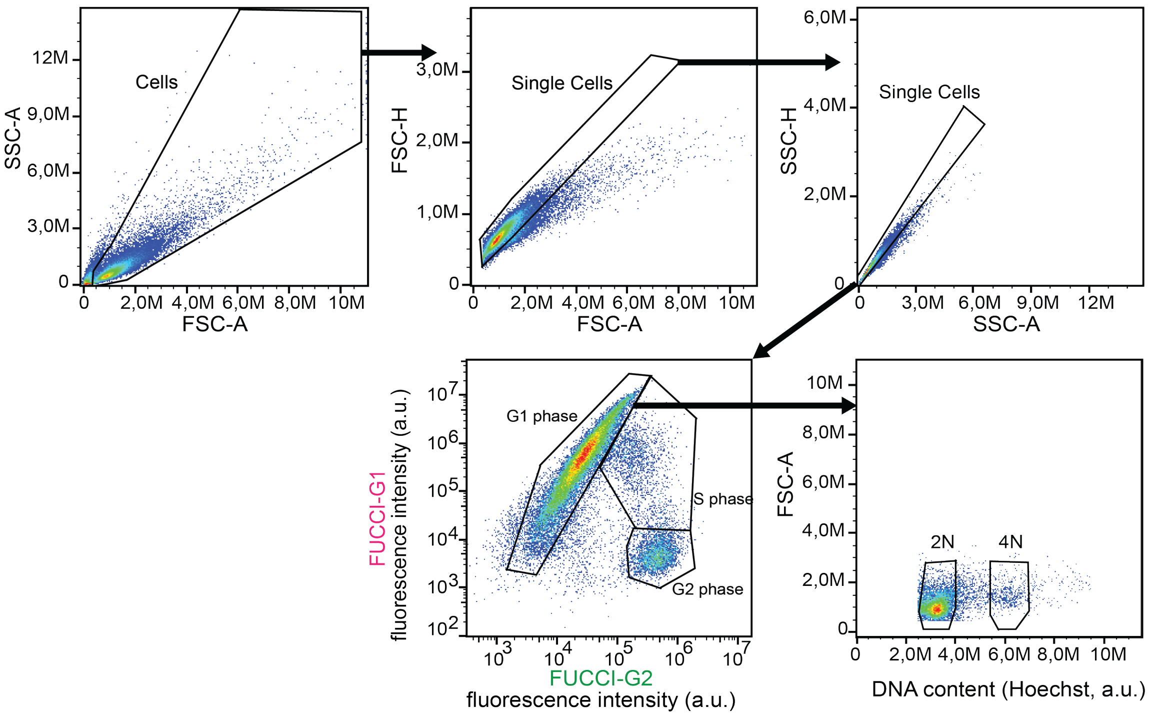


**Figure S2. Sequential gating strategy to gate single cells and G1 phase cells with 2N or 4N ploidy**

Cells were first gated based on forward scatter (FSC-A) and side scatter (SSC-A) to exclude debris and select intact cells. Doublets were then excluded through sequential gating using FSC-A versus FSC-H and SSC-A versus SSC-H to isolate single cells. G1 phase cells were identified based on FUCCI Cdt1 fluorescence intensity (high Cdt1, low Geminin). Finally, DNA content was measured using Hoechst staining to distinguish cells with 2N and 4N ploidy within the G1 population. Plots are representative of independent experiments.

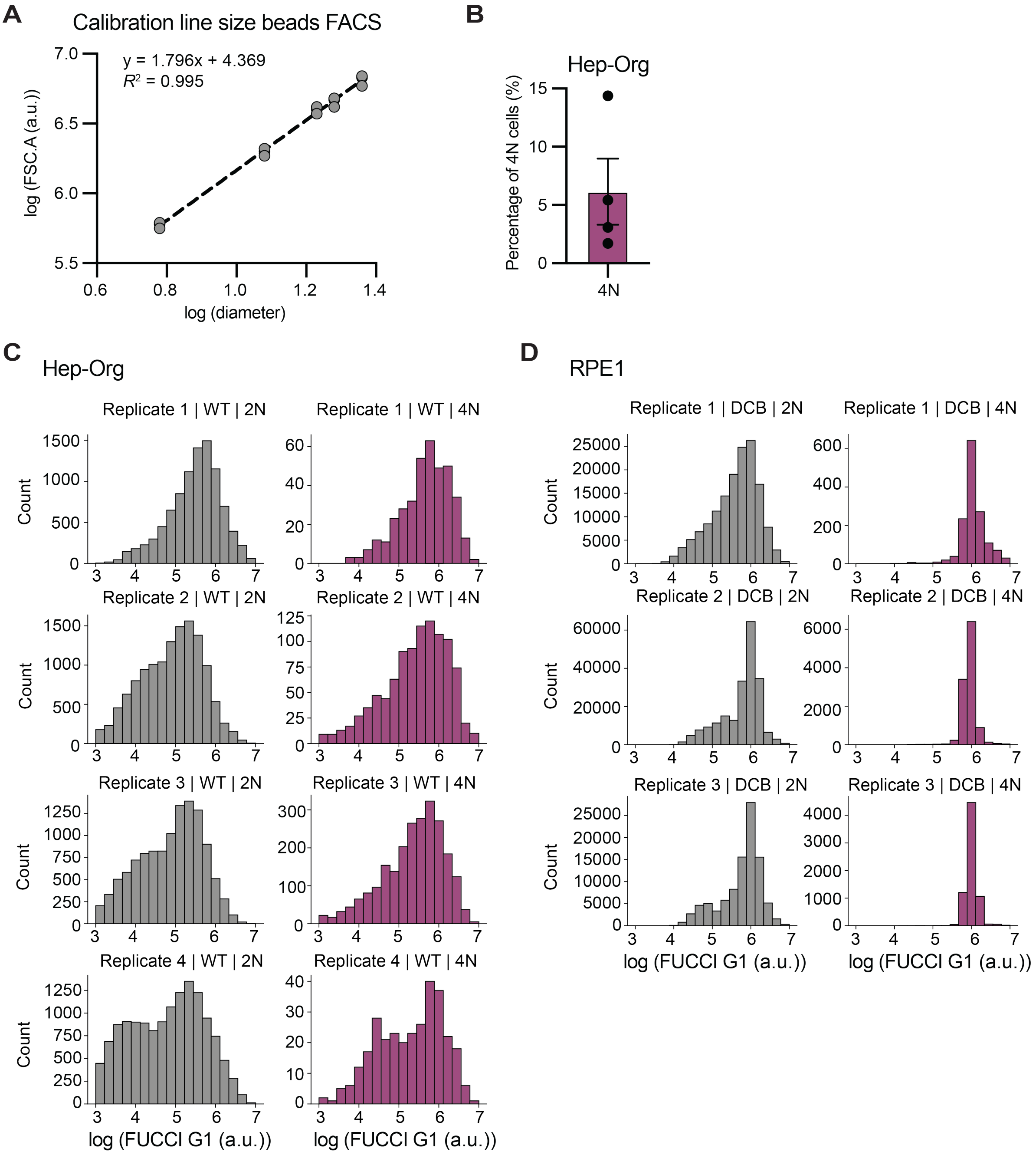


**Figure S3. Calibrated flow cytometry to measure cell size, DNA content and cell-cycle phase in RPE1 and Hep-Orgs expressing FUCCI markers**

**(A)** Calibration of flow cytometry forward scatter (FSC-A) to bead size that was used to infer cell volume. Beads with known diameters of 6, 12, 17, 19 and 23 μm were measured on the CytoFLEX that was used to perform all other flow cytometry experiments. log(diameter) was plotted against log(FSC-A) and a linear regression ($R^2$ = 0.995) was used to infer cell volume from FSC-A measurement of each cell (N = 3 independent replicates). **(B)** Percentage of binucleated cells in Hep-Orgs expressing the FUCCI markers and stained with Hoechst used for Figure 2C. Bar indicates the mean ± SEM and points show independent replicates (N = 4). **(C)** Distribution of cells in G1 based on log(Cdt1) signal in FUCCI expressing Hep Orgs. Histograms (18 bins) show similar distributions for 2N (gray, left) and 4N (purple, right) cells across N = 4 independent biological replicates. **(D)** Distribution of cells in G1 based on log(Cdt1) signal in FUCCI expressing RPE1 cells. Histograms (18 bins) show similar distributions for 2N (gray, left) and 4N (purple, right) cells across N = 3 independent replicates.

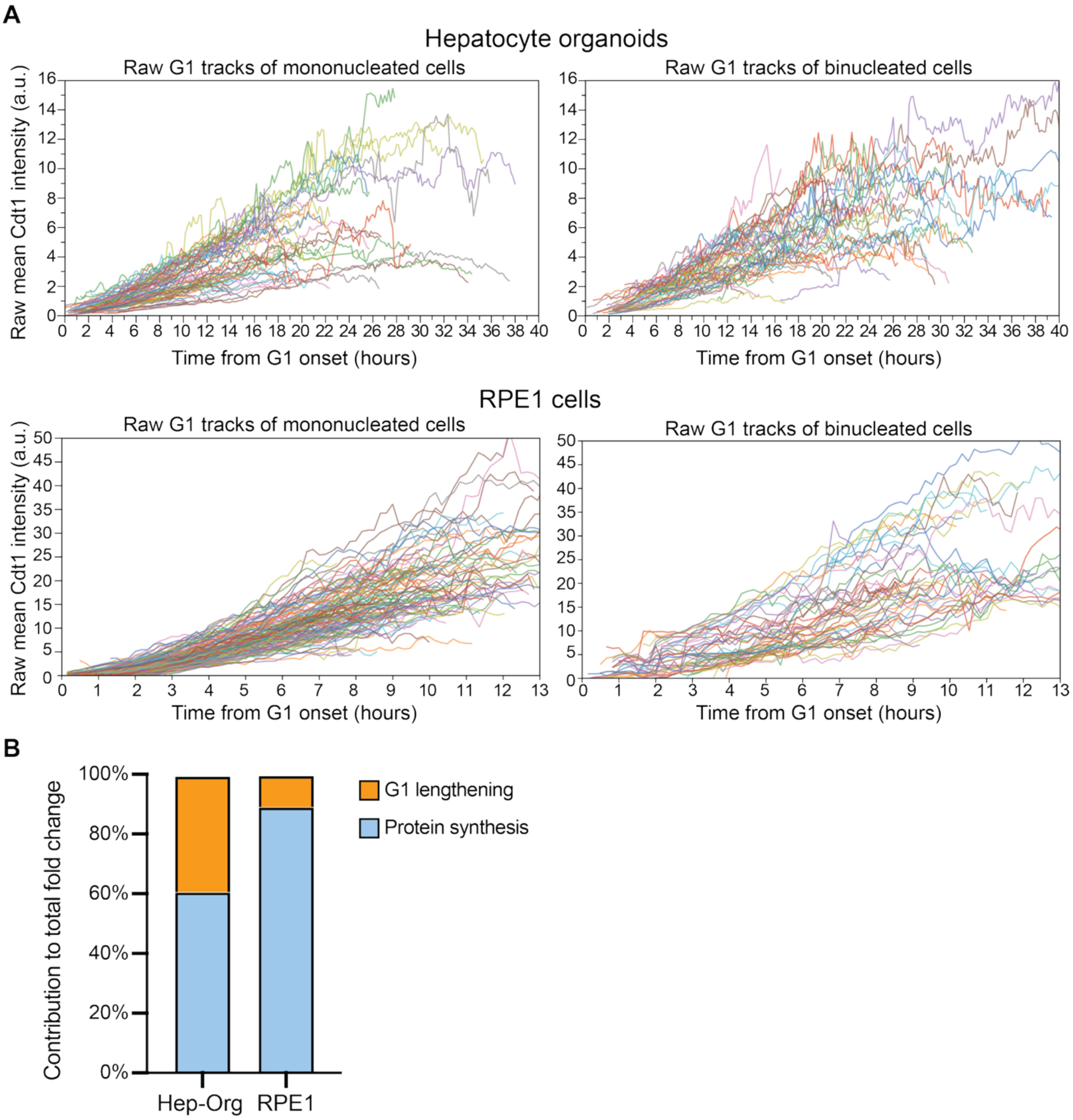


**Figure S4. Quantification of FUCCI markers by live imaging.**

**(A)** Raw tracks of Cdt1 fluorescence intensity plotted against time from G1 onset (determined as the first frame after NEB) in Hep Orgs (top) and RPE1 cells (bottom). Each line depicts the nuclear intensity of one nucleus in either mononucleated cells (left, n = 81 for Hep-Orgs and n = 130 for RPE1 cells) or binucleated cells (right, n = 50 for Hep-Orgs and n = 52 for RPE1 cells). Data is from N = 4 or N = 3 independent live-imaging experiments of Hep-Orgs and RPE1, respectively, imaged with 10-minute time intervals. **(B)** Relative contribution of increased protein synthesis rate (blue) and G1 lengthening (orange) to total protein production scaling in polyploid cells, highlighting a larger contribution of G1 extension in naturally occurring polyploid cells.

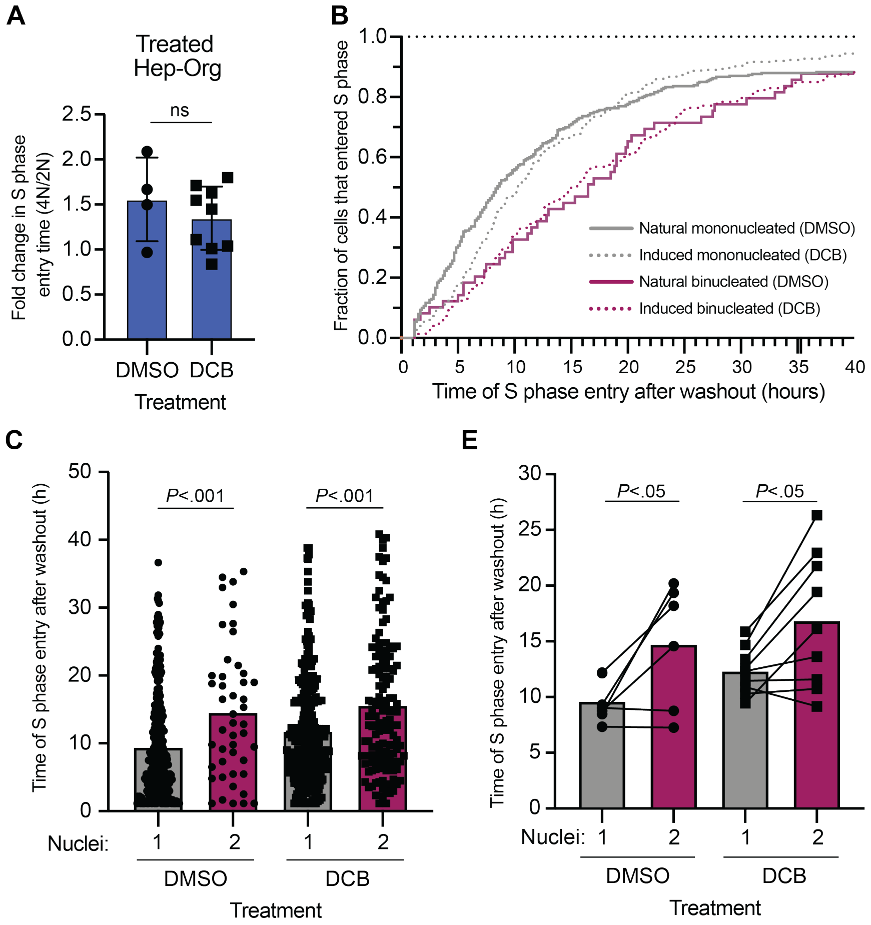


**Figure S5. Timing of S-phase entry is comparable between DMSO- and DCB-treated Hep-Orgs.**

**(A)** Fold change in time to S-phase entry in binucleated cells relative to mononucleated cells in DMSO- and DCB-treated Hep-Orgs. Each point represents the fold change in one organoid, and the bar depicts the mean ± SD. Statistical comparison was performed using a Mann-Whitney test (ns = not significant). **(B)** Cumulative fraction of cells entering S phase after washout in DMSO- and DCB-treated Hep-Orgs. Mononucleated cells (gray) enter S phase earlier than binucleated cells (purple), this delay is similar between control (DMSO, solid lines) and induced (DCB, dashed lines) polyploid cells. **(C)** Time of S-phase entry in mononucleated (gray) and binucleated (purple) Hep-Org cells following DMSO or DCB treatment. Same data as in Figure 5E, but showing individual measurements per cell. Data is from N = 2 independent experiments. Bars indicate mean and statistical comparisons were performed using a Mann–Whitney test. **(D)** Time of S phase entry following washout in DMSO- and DCB-treated Hep-Orgs. Same data as in **(C),** but now each dot represents the mean duration of either mononucleated (gray) or binucleated (purple) cells per organoid, showing S-phase entry is longer both in induced and naturally occurring polyploid cells. Lines indicate matched measurements from the same organoid and data is from N = 2 independent experiments. Bars indicate mean and statistical comparisons were performed using a paired, two-tailed Students *t*-test.

**Table S1**

| Strain | Genotype | Reference |
|---|---|---|
| GAL232 | *mcm-4(mat106 [AID::mcm-4]) I; ieSi61 [ges-1p::Tir1::mRuby + Cbr-unc-119(+)] II; unc-119(ed3) III; mcm-7(mat138 [AID::mcm-7]) V; matIs159 [pmyo-3::mCherry-PH, Pges-1::GFP-PH, Plin-48::TdTomato]* | This study |
| GAL316 | *ieSi61 [ges-1p::Tir1::mRuby + Cbr-unc-119(+)] II; unc-119(ed3) III; matIs159 [pmyo-3::mCherry-PH, Pges-1::GFP-PH, Plin-48::TdTomato]* | This study |

**Table S2**

| | | | | |
|---|---|---|---|---|
| Hep-Org Culture Medium | Advanced DMEM/F12 | 12634 | Gibco | |
| | HEPES | 15630 | Gibco | 10 mM |
| | GlutaMAX | 35050 | Gibco | 1x |
| | Penicillin/Streptomycin | 15140 | Gibco | 100 U/mL |
| | RSPO1 conditioned medium | - | (Made in house) | 15% |
| | B27 minus vitamin A | 12587-010 | Thermo Scientific | 1x |
| | Nicotinamide | 72340 | Sigma Aldrich | 10 mM |
| | N-acetylcysteine | A7250 | Sigma Aldrich | 1.25 mM |
| | Y-27632 | Y0503 | Sigma Aldrich | 5 μM |
| | CHIR99021 | 4423 | Tocris | 3 μM |
| | HGF | 100-39 | Peprotech | 50 ng/mL |
| | FGF7 | 100-19 | Peprotech | 100 ng/mL |
| | FGF10 | 100-26 | Peprotech | 100 ng/mL |
| | A83-01 | 2939 | Tocris | 2 μM |
| | EGF | AF-100-15 | Peprotech | 50 ng/mL |
| | TGF-α | 100-16A | Peprotech | 20 ng/mL |
| | Gastrin I | 3006 | Tocris | 10 nM |
